# Cortical RORβ is required for layer 4 transcriptional identity and barrel integrity

**DOI:** 10.1101/782995

**Authors:** Erin A. Clark, Michael Rutlin, Lucia Capano, Samuel Aviles, Jordan R. Saadon, Praveen Taneja, Qiyu Zhang, James Bullis, Timothy Lauer, Emma Myers, Anton Schulmann, Douglas Forrest, Sacha Nelson

## Abstract

Retinoic Acid-Related Orphan Receptor Beta (RORβ) is a transcription factor (TF) and marker of layer 4 (L4) neurons, which are distinctive both in transcriptional identity and the ability to form aggregates such as barrels in rodent somatosensory cortex. However, the relationship between transcriptional identity and L4 cytoarchitecture is largely unknown. We find RORβ is required in the cortex for L4 aggregation into barrels and thalamocortical afferent (TCA) segregation. Interestingly, barrel organization also degrades with age. Loss of RORβ delays excitatory input and disrupts gene expression and chromatin accessibility, with downregulation of L4 and upregulation of L5 genes, suggesting a shift in cellular identity. Expression and binding site accessibility change for many other TFs, including closure of neurodevelopmental TF binding sites and increased expression and binding capacity of activity-regulated TFs. Lastly, a putative target of RORβ, *Thsd7a*, is downregulated without RORβ, and *Thsd7a* knockout alone disrupts TCA organization in adult barrels.

## Introduction

Localization of function is a fundamental principle organizing mammalian brain circuitry. Structure to function mapping is particularly striking in the sensory input to L4 of the neocortex (Woolsey and Van der Loos, 1970; Catania and Kaas, 1995). L4 neurons are distinctive in their propensity to form cellular aggregates, or modules, that receive segregated thalamic inputs and represent features of the sensory periphery. Whisker barrels in the rodent somatosensory cortex are a prototypical example, but other somatosensory modules within L4 are also present in the cortices of insectivores, carnivores and primates (Krubitzer and Seelke, 2012), and columns receiving segregated input are present in the visual cortices of carnivores and primates, and in other cortical regions (Mountcastle, 1997). At the same time, gene expression studies in mouse and human show that L4 neurons also have a distinctive transcriptional identity that includes expression of RORβ (Zeng et al., 2012). Despite these two striking features, little is known about the relationships between transcriptional identity, the mechanisms that establish and regulate that identity, and features of L4 cytoarchitecture.

Researchers have long used the rodent whisker pathway to study cytoarchitecture development (Fox, 1992; Yang et al., 2018). The whisker map is organized into microcolumnar units called barrels located in primary somatosensory cortex (S1). In mice, layer 4 (L4) cortical neurons assemble into columns that form barrel walls and input is relayed via thalamocortical afferents (TCAs), which cluster in the center of barrel hollows. Each whisker is projected through corollary maps in the brainstem and ventrobasal thalamus (Van der Loos, 1976) before reaching S1.

Many proteins and pathways are required for presynaptic organization of TCAs and/or postsynaptic organization in L4 (Li and Crair, 2011; Wu et al., 2011; Erzurumlu and Gaspar, 2012). Much of what we know has focused on the requirement of input activity and intact signaling pathways. Genetic disruption of synaptic transmission via glutamate (Iwasato et al., 1997; 2000; Hannan et al., 2001; Datwani et al., 2002; Li et al., 2013; Ballester Rosado et al., 2016), or serotonin pathways (Cases et al., 1995; Salichon et al., 2001) perturb some aspect of barrel organization. Several related signal transduction pathways are also required (Abdel-Majid et al., 1998; Barnett et al., 2006; Inan et al., 2006; Watson et al., 2006; Lush et al., 2008).

Barrel formation is also regulated transcriptionally. Transcription factors (TFs) such as Bhlhe22/Bhlhb5 and Eomes are involved in the early stages of cortical arealization and barrel development (Joshi et al., 2008; Elsen et al., 2013). Downstream of these early developmental processes activity-dependent TFs, including Lmo4, NeuroD2, and Btbd3 regulate aspects of barrel organization in response to TCA inputs (Ince-Dunn et al., 2006; Kashani et al., 2006; Huang et al., 2009; Matsui et al., 2013; Wang et al., 2017). In addition, the TFs retinoic acid-related orphan receptor alpha (RORα) and beta (RORβ), are also implicated in barrel formation. RORα and RORβ are expressed in regions of the somatosensory barrel map, with RORα expressed in brainstem, thalamus and cortex, and RORβ in thalamus and cortex (Nakagawa and O’Leary, 2003). Recently, RORα was shown to be required in the thalamus and cortex for proper TCA segregation and barrel wall formation (Vitalis et al., 2017). Mis-expression of RORβ in neocortex is sufficient to drive cortical neuron clustering and TCA recruitment to ectopic barrel-like structures (Jabaudon et al., 2012). Together these studies have identified multiple TFs with major roles in early barrel development that likely set the stage for more downstream terminal differentiation TFs and activity-regulated TFs to hone the network. Early cortical development, TCA pathfinding, and activity dependent gene regulation are prolific areas of research. However, the later stages of neuronal specification and the molecular mechanisms of TFs involved in barrel development are currently underexplored. TFs such as Bhlh5 and Eomes have broad roles and are widely expressed in the cortex while the more narrowly expressed TFs such as Btbd3 are downstream of activity input leaving a gap in our understanding of the intermediate steps that connect cortical development to activity driven processes. Given the restricted layer specific expression of RORβ and its upregulation concomitant with the final stages of barrel formation and the onset activity input, we hypothesized it would be a good candidate to study transcriptional mechanisms connecting cellular specification in L4 with cytoarchitecture and network development.

We show that in addition to being sufficient, RORβ is also required for both pre- and postsynaptic barrel organization. Without RORβ in the cortex, L4 neurons fail to migrate tangentially and organize into barrel wall structures. This also reduced TCA segregation shortly after barrel formation would have normally occurred. Interestingly, TCA segregation also declined as animals aged. Without RORβ, L4 gene expression and chromatin accessibility were disrupted, with L4-specific genes downregulated and L5-specific genes upregulated suggesting a shift in terminal cellular identity. This involved complex changes in the expression and/or chromatin accessibility at binding motifs for many TFs in addition to RORβ, including developmental regulators and activity-regulated TFs. L4 neurons also received delayed excitatory input, a key step in barrel development. Lastly, we identify a putative direct gene target of RORβ, *Thsd7a*, that is downregulated without RORβ and is required for maintained TCA organization in adulthood. Together these data characterize the role of RORβ across multiple levels to connect molecular and transcriptional mechanisms to cortical organization and place RORβ as a key regulator of a complex developmental transition orchestrating terminal L4 specification and initiating activity responsiveness.

## Results

Cortical barrels in mice are complex structures. Cell-sparse barrel hollows are where thalamic projections are concentrated. Barrel walls are formed by cortical cell aggregates that surround the thalamocortical afferents (TCAs). Barrel septa consist of the intermediate spaces between barrel walls (Woolsey and Van der Loos, 1970). To assess the impact of RORβ loss on barrel organization we used two staining methods. Barrel walls were visualized by Nissl staining (Van der Loos and Woolsey, 1973) and barrel hollows were visualized by vesicular glutamate transporter 2 (VGLUT2), which is strongly expressed in TCAs (Fremeau et al., 2001; Liguz-Lecznar and Skangiel-Kramska, 2007; Liu et al., 2013). This strategy allowed clear identification of changes in either structure independently. Cytochrome oxidase (CO) staining was also used in some conditions, but the presence of CO signal in both barrel walls and TCAs made it less useful.

### RORβ is required for postnatal barrel wall formation and influences segregation of thalamocortical afferents (TCAs)

To begin exploring RORβ function in barrel organization, we used a global, constitutive knock-out (KO), which contains a GFP expression cassette knocked into the *Rorb* locus. *Rorb*^*GFP/+*^ mice express GFP in RORβ expressing cells allowing identification of barrel cortex without significant disruption to barrel structures or neuronal function (Rice and Van der Loos, 1977). *Rorb*^*GFP/+*^ mice were used as controls (Ctl), while *Rorb*^*GFP/GFP*^ mice disrupt both copies of *Rorb* to generate a KO. Controls showed no detectable disruption to barrel organization compared to WT animals (compare Figures 1A, 2A to Figure 7C).

**Figure 1.**
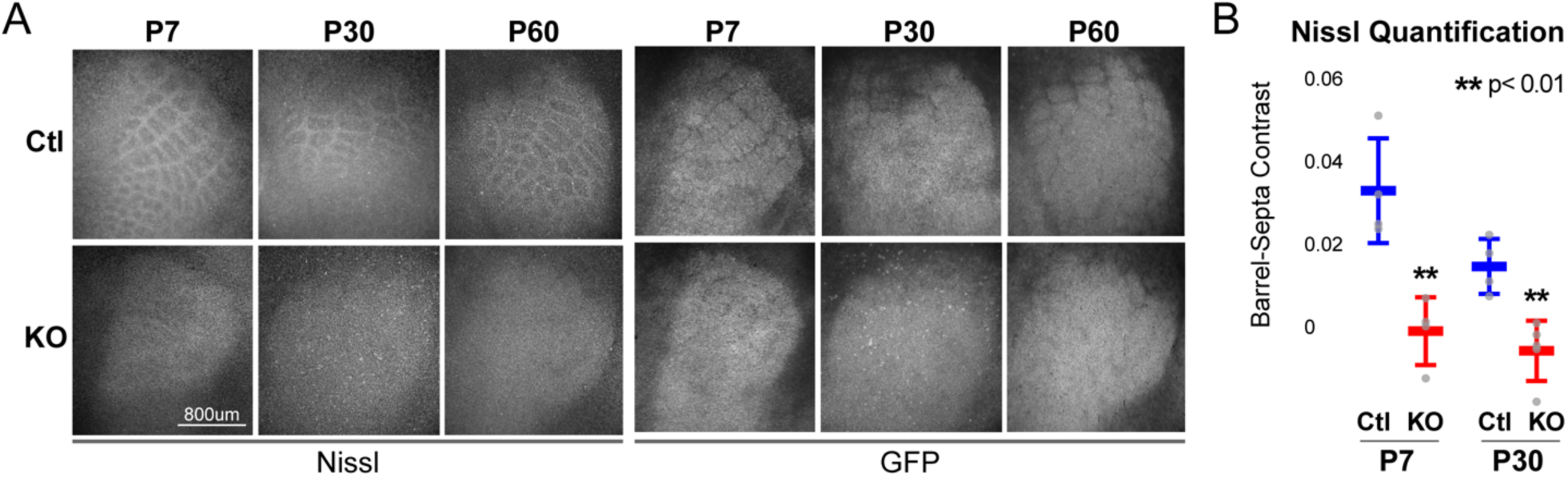
RORβ is required for postnatal barrel wall formation. Nissl staining on tangential sections of flatten cortices after global, constitutive knock-out shows barrel wall organization requires RORβ. (A) Top two rows show Nissl staining in whisker barrel field as identified by strong GFP expression (bottom two rows). Control (Ctl) and *Rorb* knock-out (KO) animals were age matched at P7, P30, and P60. (B) Quantification of barrel hollow to barrel walls/septa contrast (Barrel-Septa Contrast) from Nissl staining. N=4 age-matched animals for each genotype (Ctl or KO). Two tissue sections containing the largest portions of whisker barrel field identified by GFP signal were averaged per animal. Whisker plots show the median per animal ± standard deviation. Gray points show mean contrast for each animal. P-value by independent sample t-test, between Ctl and KO at each timepoint.

**Figure 2.**
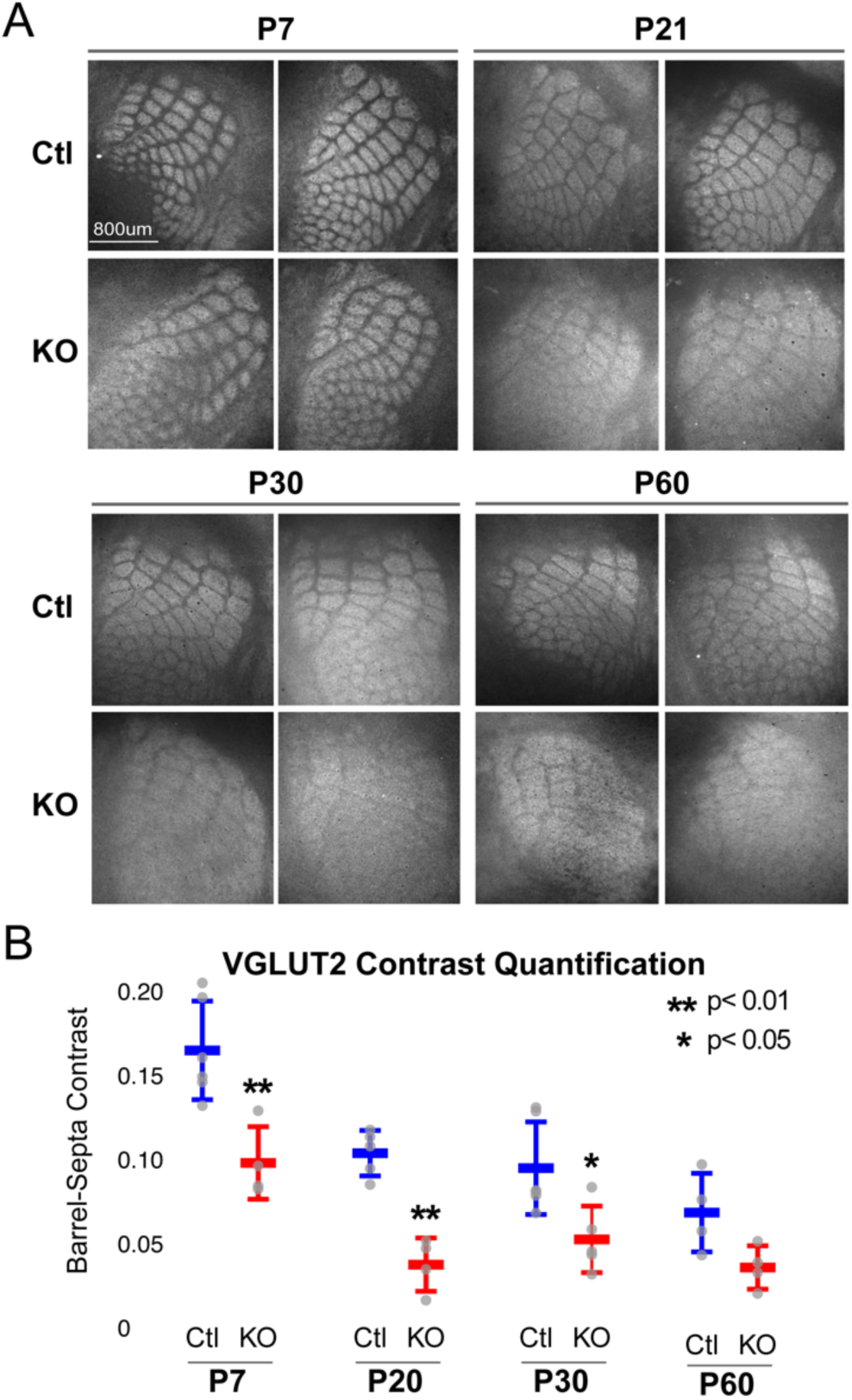
*Rorb* KO reduces thalamocortical afferent (TCA) segregation. (A) VGLUT2 staining of excitatory thalamic axon terminals in cortical whisker barrels shows normal initial TCA patterning at P7 but degradation of barrel-septa contrast in *Rorb* KO adults. Ctl and *Rorb* KO animals were age matched. (B) Quantification of barrel hollow to barrel walls/septa contrast (Barrel-Septa Contrast) in VGLUT2. N = 4-6 agematched animals for each genotype (Ctl or KO). Two tissue sections containing the largest portions of whisker barrel field identified by GFP signal were averaged per animal. Whisker plots show median contrast per animal ± standard deviation. Gray points show mean contrast for each animal. P-value by independent sample t-test, between Ctl and KO at each timepoint.

Barrels form around postnatal day 5 (Oishi et al., 2016a). Nissl staining of barrel walls at P7, P30, and P60 showed that RORβ is required for barrel wall formation. Representative images of Nissl and GFP are shown in Figure 1A where the lack of barrel wall organization is clearly visible at P7 and remains disrupted at P30. Figure 1B quantifies this effect as the contrast between barrel hollows and barrel wall/septa fluorescence intensity. Contrast was calculated as (barrel - septa) / (barrel + septa) where septa includes barrel walls (see methods for details). Quantification demonstrated a near complete lack of contrast in KO barrel cortex supporting a lack of cortical organization.

While TCAs have been shown to instruct cortical cell organization we hypothesized the lack of barrel walls might reciprocally affect TCA organization. TCAs visualized by VGLUT2 staining showed an intact pattern of barrel hollows at P7 in KO animals, Figure 2A. However, careful quantification of the VGLUT2 contrast between hollows and septa showed a significant decrease in the KO suggesting loss of RORβ and/or the lack of barrel walls had a mild but measurable effect on TCA segregation. Interestingly, as animals aged into adulthood TCA segregation also declined in control as well as *Rorb* KO animals. Disorganization in the *Rorb* KO was characterized by both loss of quantifiable VGLUT2 contrast as well as the qualitative barrel patterning most obvious at P60 between Ctl and KO in Figure 2A. Both genotype and age significantly affected VGLUT2 contrast (genotype p=4.5e-07 and age p=2.6e-06 by two-way ANOVA) but did not interact significantly. This suggests that while both age and loss of RORβ significantly reduced contrast, loss of RORβ did not significantly change the time course of TCA desegregation. Together these data show that RORβ is critical for normal whisker barrel formation and, loss of TCA segregation into adulthood suggests time/age affects cytoarchitecture.

### RORβ is required in the cortex but not the thalamus for barrel organization

In addition to L4 excitatory neurons, RORβ is expressed in the thalamic neurons that project to barrel hollows. To assess whether the disruption of barrels is dependent on RORβ expression in thalamus and/or locally in cortex we used a floxed allele of *Rorb* (*Rorb*^*f/f*^) crossed to Cre-driver lines generating tissue-specific disruption of RORβ as diagrammed in Figure 3A. A knockin line expressing Cre from the serotonin transporter gene, Sert (Slc6a4 or 5-HTT) locus was used to knockout *Rorb* in the thalamus. The Sert-cre line alone showed a mild disruption to TCA organization without disrupting barrel walls, suggesting the Cre knockin might be hypomorphic (Figure 3B-C). However, thalamic KO of *Rorb* (Sert-cre; *Rorb*^*f/f*^) showed no additional disruption to TCAs or barrel walls. Thus, loss of RORβ in thalamic neurons was not responsible for the loss of cortical wall organization or the majority of TCA disorganization observed in the global *Rorb*^*GFP/GFP*^ KO.

**Figure 3.**
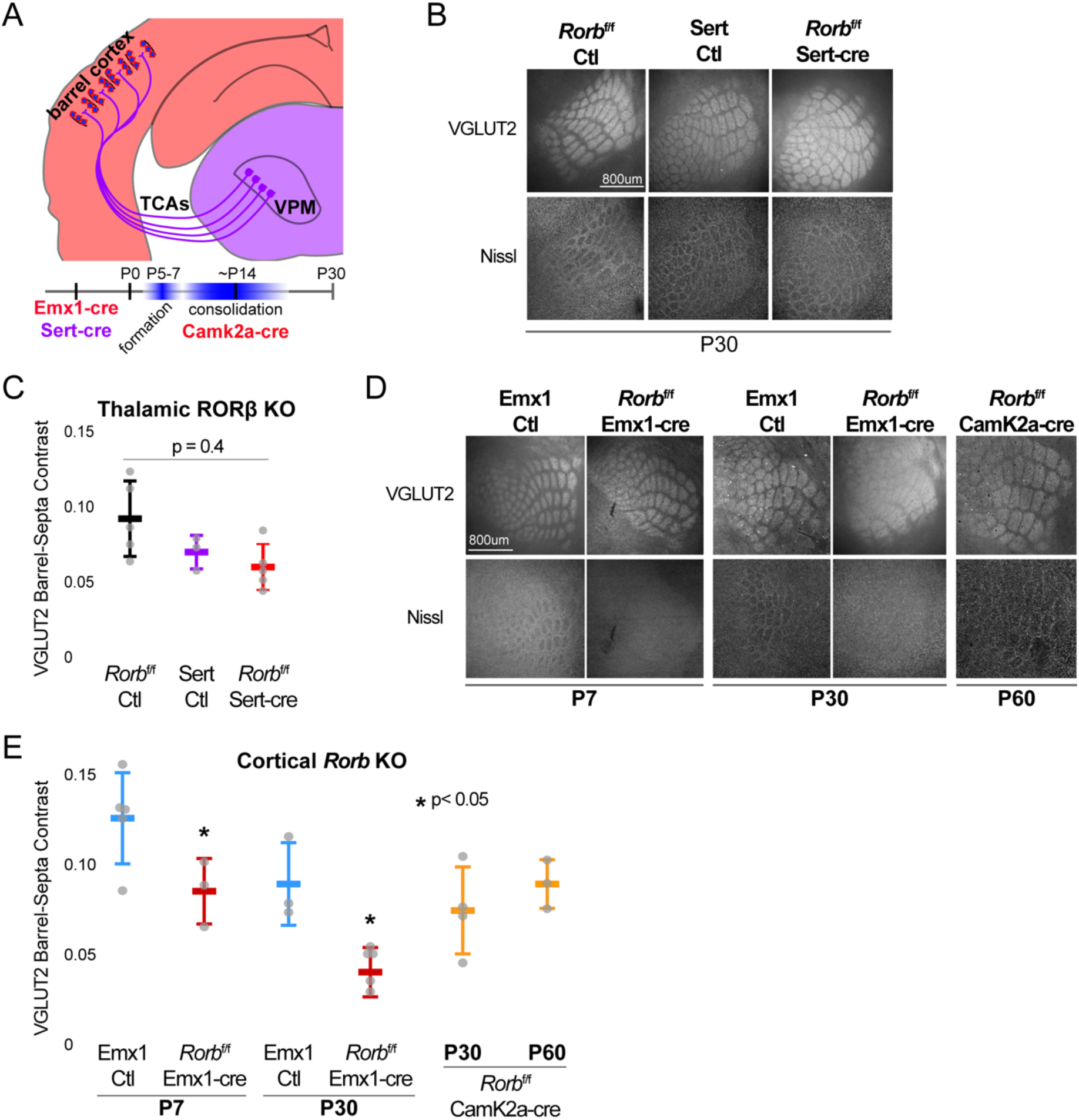
RORβ is required in the cortex but not the thalamus for barrel organization. (A) Diagram and timeline of Cre driver line tissuespecific expression in cortex versus thalamus and timing relative to barrel formation and consolidation. Color indicates expression in cortex (red) or thalamus (purple). (B) VGLUT2 and Nissl staining of whisker barrel cortex at P30 from floxed *Rorb* control without Cre (*Rorb*^f/f^ Ctl), Sert-cre control (Sert Ctl) without floxed *Rorb* and the cross (*Rorb*^f/f^ Sert-cre), which knocks out RORβ specifically in thalamus during embryonic development. thalamus during embryonic development. Whisker plots as described for Figure 1B. (C) Quantification of VGLUT2 Barrel-Septa Contrast in genetic lines from B. N=3-5 P30 animals. Quantification and plotting as described in Figure 2B. P-value by ANOVA. (D) VGLUT2 and Nissl staining of whisker barrel cortex from Emx1-cre control (Emx1-cre Ctl) without floxed *Rorb*, and the cross (*Rorb*^f/f^ Emx1-cre) from P7 and P30 animals, and a P60 animal from floxed *Rorb* crossed to a CamK2a-cre driver line. Emx1-cre knocks out RORβ specifically in forebrain during embryonic development, and CamK2a-cre knocks out RORβ in forebrain neurons at postnatal weeks 2-3. (E) Quantification of VGLUT2 Barrel-Septa Contrast in genetic lines from D. N=3-5 animals per age group. Quantification and plotting as described in Figure 2B. P-values by independent sample t-test, between Emx1 Ctl and KO at each time point. Whisker plots as described for Figure 1B.

A knock-in line expressing Cre from the Emx1 locus removed RORβ specifically in forebrain structures. Emx1-cre alone showed no significant disruption to barrel organization (Figure 3D-E). However, barrel organization was significantly disrupted by cortical KO of *Rorb* (Emx1-cre; *Rorb*^*f/f*^). In addition, a CamK2a-cre diver line that removes RORβ in the cortex after barrel formation, showed no effect. Together these data demonstrate that RORβ is required in the cortex prior to barrel formation. Loss of RORβ in thalamus or after barrels have formed does not disrupt barrel architecture, suggesting RORβ drives barrel wall organization through cell-intrinsic mechanisms within layer 4.

### RORβ is required for expression of a layer 4 gene profile and repression of layer 5 genes

Because RORβ is a transcription factor we hypothesized loss of function would change gene expression in L4 neurons. To test this, RNA-seq was performed on sorted GFP^+^ cells from micro-dissected L4 S1. We were careful in this dissection to exclude a small population of GFP^+^ L5 neurons. Differential expression analysis between *Rorb*^*GFP/+*^ and *Rorb*^*GFP* /GFP^ cells identified many dysregulated genes (fold change ≥ 2, adjusted p-value < 0.01). At postnatal day 2 (P2) and prior to barrel formation, 246 genes were significantly disrupted with 51% downregulated in the KO. At P7, just after barrel formation, 433 genes were disrupted with 36% downregulated. At P30, 286 genes were disrupted with 37% downregulated. Examining the overlap between ages we find very few genes significantly disrupted in the same direction across time points, suggesting highly dynamic and complex regulation, Figure 4-figure supplement A.

RORβ expression is a key feature distinguishing L4 neurons (Lein et al., 2007). To examine the effect of RORβ loss on layer-specific identity we assessed the layer specificity of differentially expressed genes (DEGs) using the Allen Brain Atlas (Doyle et al., 2008). Genes were considered layer-specific if the *in-situ* hybridization (ISH) signal appeared at least three-fold higher in one layer (considering layers 2 and 3 together). Many genes had complex specificities showing enrichment in two or more layers. These were not included for simplicity. Grouping DEGs based on their normal layer-specific expression pattern we see overall downregulation of superficial layer genes with a modest effect on layer 2/3 genes and stronger loss of L4 gene expression in the KO, Figure 4A-B. In addition, deep layer genes were generally upregulated in the KO with the strongest effect on layer 5-specific genes. Several L5 genes are worth noting. Bcl11B/Ctip2, is a marker of thick-tufted L5B-type neurons and significantly upregulated at P2 in the KO, but silenced at P7 and P30 similar to control (Figure 4-figure supplement B). Fezf2, another L5B marker and regulator of *Bcl11B*, was similarly silenced over barrel development, but was mildly overexpressed at P30 in the KO. Foxo1, is mainly expressed in L5 at younger ages (Allen Developing Mouse Brain Atlas) and shows a decline in expression over barrel development but, was significantly overexpressed in the KO at P7. *ETV1*, also a L5A marker (Doyle et al., 2008), was upregulated in the KO at both P2 and P30. Lastly, *EGR4* was upregulated at P30 in the KO, and has been associated with ETV1 expressing neurons (Buenrostro et al., 2015). Together these data support a general but disorganized shift in layer identity with many different factors implicated at distinct time points.

**Figure 4.**
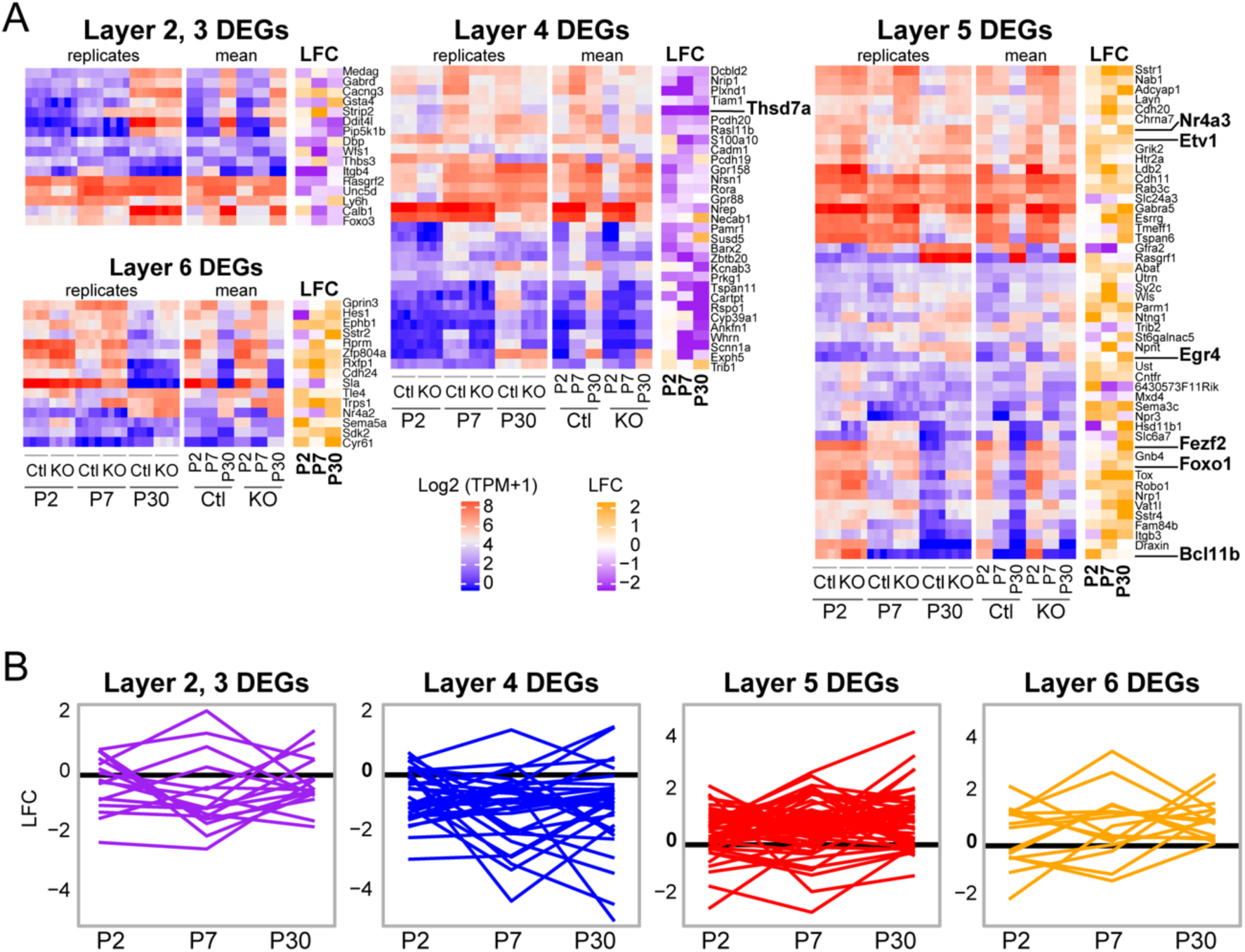
*Rorb* KO shifts the expression profile of neurons from a layer 4 to layer 5. (A) Heatmaps showing marker genes or genes strongly enriched, as identified in the Allen Brain Atlas, for each layer of the neocortex. Log-transformed TPMs are color scaled in red and blue for each of the four RNA-seq replicates in the left most heatmap and the mean for each time point and genotype in the middle heatmaps. Log fold change (LFC) between control (Ctl) and *Rorb* KO is color-scaled in orange and purple in the right most heatmaps. (B) Line plots showing LFC for the same genes. The black line indicates no change. Negative LFC indicated decreased expression in *Rorb* KO, and LFC > 0 indicate increased expression in *Rorb* KO.

### *Rorb* KO disrupts transcription factor binding sites near DEGs

RORβ, Bcl11b, Foxo1, Etv1, and Egr4 are TFs that often regulate gene expression by binding to distal regulatory sites such as enhancers. There are many chromatin features of enhancers, one of which is that they are open and accessible to enzymatic digestion in assays such as the Assay for Transposase Accessible Chromatin (ATAC) (Chen et al., 2014). To begin examining mechanisms involved in changing gene expression, we performed ATAC-seq on sorted GFP^+^ L4 neurons from control and *Rorb* KO animals at P30 (Figure 5A). High confidence ATAC-seq peaks were assessed for differential accessibility between control and KO samples. We identified 5,210 peaks with ≥ 2-fold change in accessibility (FDR < 0.02). Nearly 4-times as many regions lost accessibility (N=4,123 closed) than increased (N=1,087 opened), Figure 5-figure supplement A. Differential ATAC peaks were primarily located in introns and intergenic regions (Figure 5-figure supplement B) suggesting loss of RORβ function resulted in closure of many more regulatory regions than opening.

**Figure 5.**
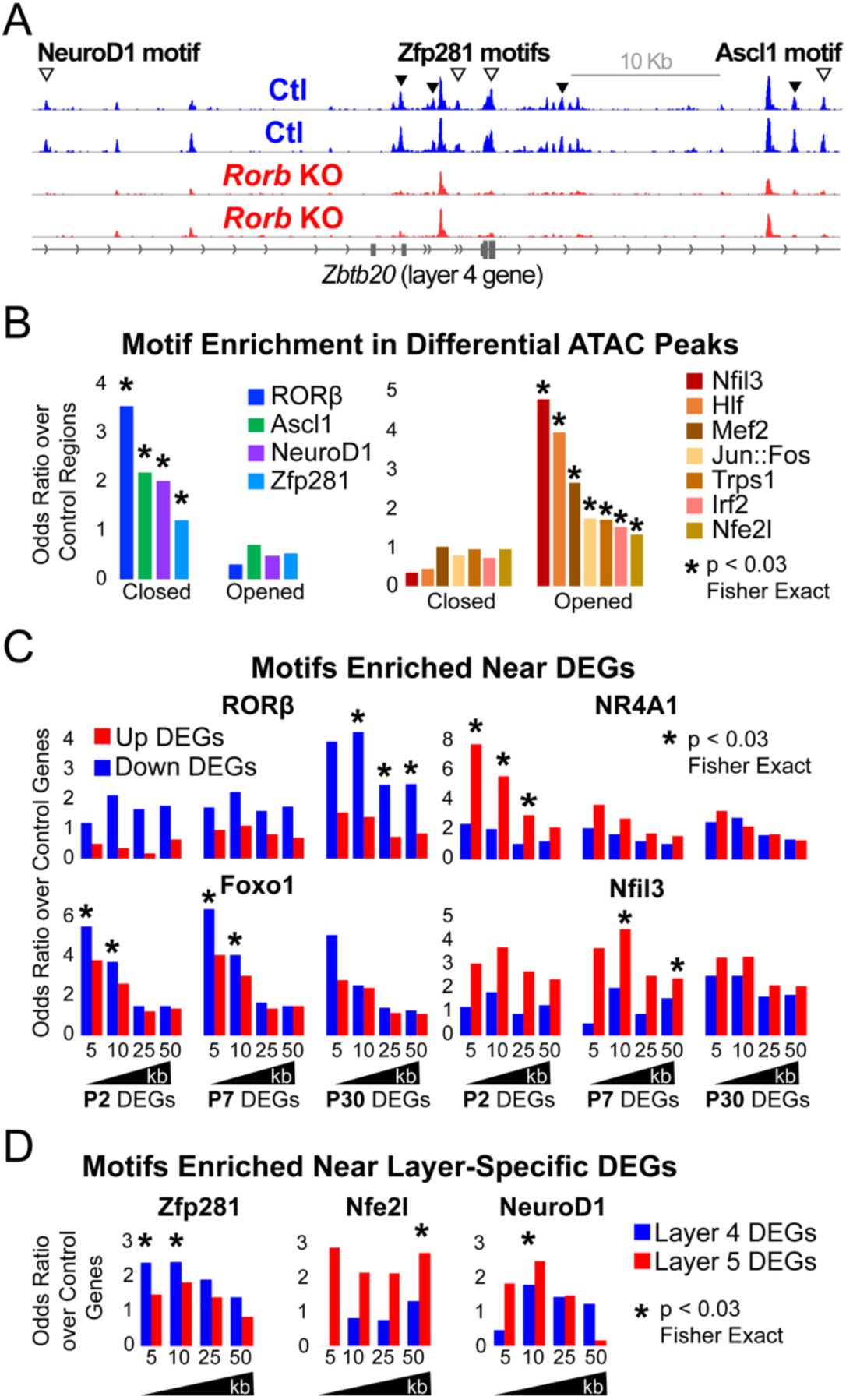
*Rorb* KO disrupts transcription factor binding sites near DEGs. (A) ATAC-seq normalized reads per million (RPM) for biological replicates. Samples collected from GFP^+^ S1 layer IV RORβ^gfp/+^ neurons (Ctl, blue) and GFP^+^ S1 layer IV RORβ^gfp/gfp^ neurons (KO, red). Arrows indicate differential peaks (fold change ≥ 2, FDR<0.02). Open arrows indicate differential peaks with transcription factor motif sequences as in (B). (B) Cross-validated motifs with significant enrichment in ATAC peaks with differential accessibility. Closed; regions with significantly reduced access, Opened; regions with significantly increased access in the *Rorb* KO. Motif instances were cross-validated between MEME and HOMER algorithms. Odds ratio and p-value calculated comparing to motif frequency in control regions. (C-D) Cross-validated motif enrichment in ATAC peaks near the TSSs of (C) upregulated or downregulated DEGs and (D) layer 4- or 5- specific genes. Bars plot odds ratio over control regions. Asterisk indicates significant motif enrichment (p<0.03 by Fisher exact test) in nearby ATAC peaks compared to control regions and separately significant enrichment (p<0.03 by Fisher exact test) of DEGs with a nearby motif compared to an independent group of control genes.

We hypothesized that many of the closed regions might contain a RORβ binding motif while regions that opened may have binding potential for other TFs. To assess this possibility, two software algorithms (MEME and HOMER) were used to identify *de novo* enriched motifs from the DNA sequences of differential ATAC peaks separating closed and opened regions. This unbiased analysis also identifies which enriched sequences match known TF binding motifs. RORβ was the top motif from closed regions, Figure 5-figure supplement C. Considering only expressed TFs, the potent neurogenic factors NeuroD1 and Ascl1 were also among the top motifs in closed regions. In regions that opened, the top motifs from expressed TFs were Nfil3, Hlf, Jun, Fos, Trps1, Mef2a/c/d and Irf2. Similar analysis was performed on ATAC peaks within 10 kb of up or downregulated DEGs as well as L4 and L5 DEGs. To confirm enrichment and identify motif locations we used MEME FIMO and HOMER to scan for instances of a given set of motifs. This was done for all expressed TFs either enriched in the *de novo* motif analysis or differentially expressed, for which high quality motif models existed. Motif instances were cross-validated by retaining only those found by both MEME and HOMER. Figure 5B plots the odds ratio of motifs significantly enriched compared to control regions. Many of the motifs found by *de novo* analysis were confirmed, including RORβ in regions that closed.

To assess which TFs might play a significant role in up or downregulation of DEGs we varied a distance window around the transcription start site (TSS) to identify nearby ATAC or control regions containing a DNA motif. We tested for enrichment of motifs in ATAC regions near DEGs compared to motifs in control regions. We also tested whether DEGs with a nearby motif were significantly enriched compared to a control group of genes that did not change expression in the *Rorb* KO. In essence, we tested whether motifs were enriched around certain DEGs and whether a significant portion of those DEGs had a nearby motif. To reduce false positives, only motifs with significant enrichment in both tests are shown in Figure 5C-D.

Genes downregulated at P30 showed significant enrichment of nearby RORβ motifs suggesting RORβ is important for gene activation (Figure 5C). Motifs for Nr4a1 and Nfil3 were enriched near upregulated DEGs at P2 and P7 respectively consistent with an early role for these TFs in activating expression. Foxo1 motifs were enriched near genes downregulated at P2 and P7. Consistent with a role in early gene regulation, Foxo1 was highly expressed at P2 and declined with age in control neurons (Figure 4-figure supplement B). However, in the KO, Foxo1 remained significantly elevated at P7 eventually decreasing to levels comparable to control at P30. The close proximity of Foxo1 binding sites to downregulated genes and its elevated expression at younger ages suggests it may act as a repressor that is normally silenced just after barrel formation to allow proper gene induction in L4 neurons. Without RORβ, silencing of Foxo1 is delayed allowing it to aberrantly repress targets at younger ages.

Interestingly, we did not find RORβ motifs enriched near L4 genes suggesting the shift in layer-specific gene expression is a downstream effect of RORβ loss. While RORβ does not appear to directly regulate layer specific genes, Zfp281 motifs were enriched near L4 genes in the *de novo* motif search and confirmed by specific mapping (Figure 5-figure supplement C and Figure 5D). Zfp281 was highly expressed in both samples, at all ages, and unchanged by *Rorb* KO (Figure 5-figure supplement D). Zfp281 motifs were also enriched in regions that closed in the *Rorb* KO suggesting it might be a novel activator of L4-specific genes and dependent on some other factor to maintain accessible chromatin at its binding sites.

Nfe2l and NeuroD1 motifs were enriched near L5 genes. NeuroD1 motifs were also enriched in regions that closed suggesting it might act as an inhibitor of L5-specific genes as these genes increased expression when NeuroD1 sites closed. Nfe2l consists of a family of TFs that share a binding motif. Nfe2l1 was expressed at younger ages and increased in the adult while Nfe2l3 was highly expressed at P2 and silenced by P7 (Figure 5-figure supplement D). *Rorb* KO did not significantly disrupt expression of either, but the motif was enriched in regions that opened suggesting Nfe2l1 and/or 3 may be novel activators of L5-specific genes.

The TF motifs enriched near upregulated DEGs were noteworthy for possible relationships with neuronal activity. Nr4a1 is an activity induced TF that regulates the density and distribution of excitatory synapses (Mitsui et al., 2001). Nfil3 and Hlf bind and compete for similar DNA motifs (Beaumont et al., 2012), and may also be involved in activity-regulated transcription. Nfil3 is upregulated in human brain tissue following seizures (Beaumont et al., 2012), and mutations in Hlf are linked to spontaneous seizures (Gachon et al., 2004; Hawkins and Kearney, 2016). In addition, motifs for the classic immediate early genes, Jun and Fos, were enriched in regions that opened. These observations led us to examine the expression of other activity-regulated TFs. Many were significantly upregulated at P30 while Lmo4 and its binding partner Lbd2 were upregulated at P7 (Figure 5-figure supplement E). Lmo4 expression is induced by calcium signaling and is required for TCA patterning in barrel cortex (Kashani et al., 2006; Huang et al., 2009). Another activity-regulated TF, Btbd3, which drives L4 neurons to orient their dendrites into barrel hollows, was significantly downregulated (Figure 5-figure supplement E). Lmo4 and Btbd3 are the only genes previously shown to disrupt barrels that were also dysregulated in the *Rorb* KO (Figure 5-figure supplement F). Interestingly, *Lmo4* KO disrupts barrels yet it was upregulated in the *Rorb* KO indicative that *Rorb* KO disrupts barrels through a divergent mechanism from what has been previously described.

Interestingly, *S100A10*, is another gene associated with L5A neurons (Schmidt et al., 2012; Svenningsson et al., 2013), but was downregulated at P7 and P30 (Figure 5-figure supplement G). The protein product of *S100A10*, p11, is involved in serotonin signaling via binding to the serotonin receptors Htr1b, Htr1d, and Htr4 (Warner-Schmidt et al., 2009). Htr1b was the only serotonin receptor expressed in our samples and was also significantly downregulated at P7 and P30. These data suggest that in addition to altered layer identity, *Rorb* KO may also disrupt serotonergic signaling, an important pathway in TCA communication with cortex (Kawasaki, 2015). Together with upregulation of activity-regulated TFs, L4 neurons in the *Rorb* KO likely have significantly altered responses to activity.

These analyses paint a complex picture where gene expression in L4 *Rorb* KO neurons is disrupted by multiple mechanisms. Loss of RORβ results in closure of many RORβ binding sites which are also enriched near genes with reduced expression in adults consistent with an activator role for RORβ. Other regulatory changes involve complex combinations of altered TF expression and/or altered binding potential at sites that opened or closed in the KO likely due to downstream effects of RORβ loss. These changes impact both known neurodevelopmental regulators as well as activity-regulated TFs.

### *Rorb* KO delays excitatory input to barrel cortex

To examine whether RORβ loss impacts network activity we examined inhibitory and excitatory synaptic properties of L4 neurons. We found no change in inhibitory innervation at P14 or P24 as measured by miniature inhibitory postsynaptic currents (mIPSCs), Figure 6-figure supplement A-B. However, synaptic function as measured by miniature excitatory postsynaptic currents (mEPSCs) revealed a significant delay in excitatory input, Figure 6A-C. At P5, shortly after thalamocortical LTP has ended (Feldman et al., 1998), the frequency of mEPSCs was low and comparable in control and KO Figure 6B-C. At P7, when recurrent cortical synapses begin to sharply increase (Ashby and Isaac, 2011), controls showed increased mEPSC frequency. However, *Rorb* KO animals had a significantly lower mEPSC frequency at P7 (Figure 6A-C), suggesting decreased functional synaptic input. At P10, *Rorb* KO neurons increased mEPSC frequency to levels comparable with controls. This suggests synaptic connections were delayed by *Rorb* KO mostly likely affecting recurrent excitatory connections. At P10, this defect in frequency is mostly corrected, but *Rorb* KO also showed significantly increased mEPSC amplitude at P10, possibly compensating for the delay at P7. These data support a subtle functional disruption to the barrel circuit in *Rorb* KO animals that is consistent with the transcriptional changes.

**Figure 6.**
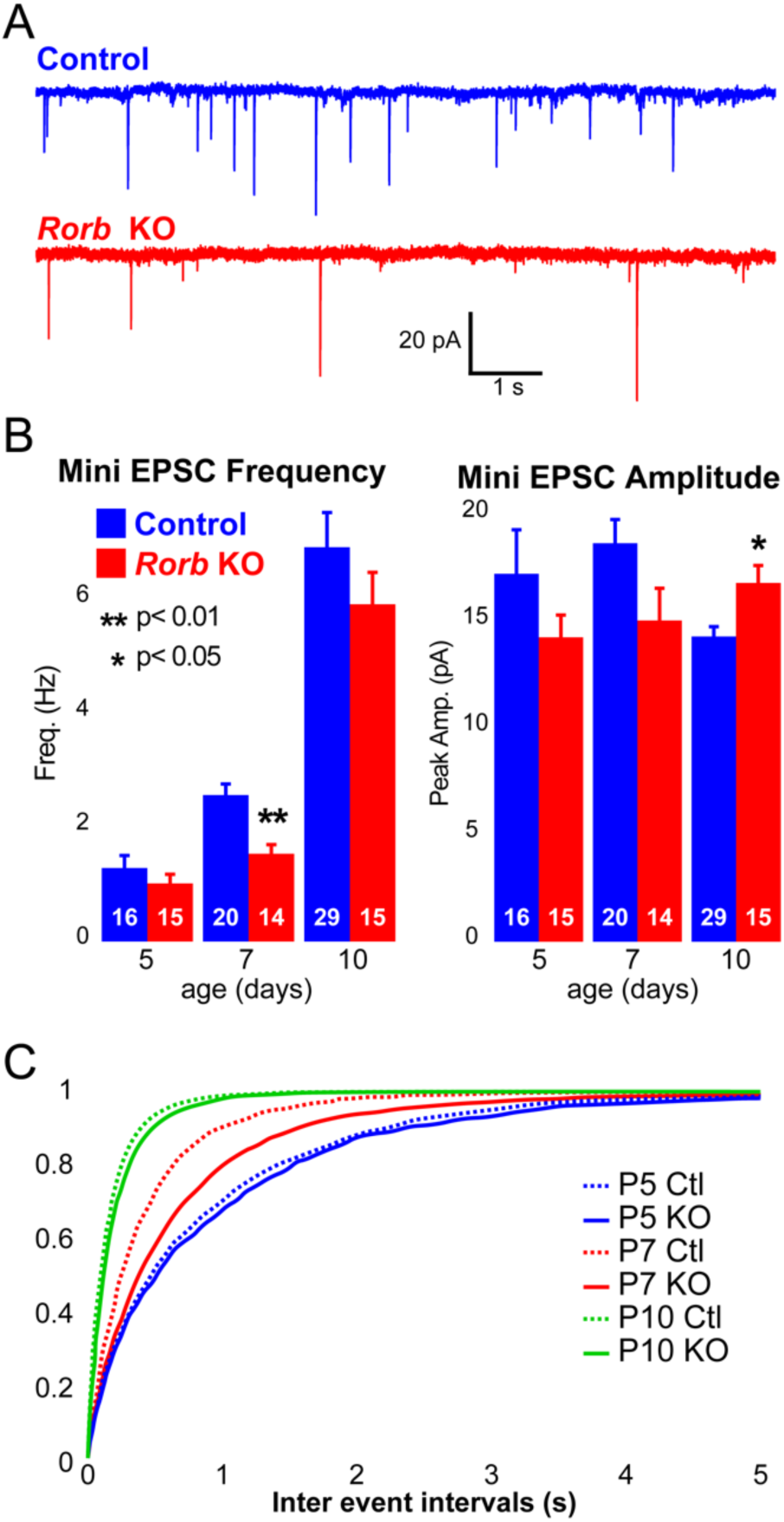
*Rorb* KO delays excitatory input to barrel cortex. (A) Example of mini excitatory postsynaptic currents (mEPSCs) from L4 barrel cortex at P7. (B) Average mEPSC frequency and from Ctl and *Rorb* KO L4 barrel cortex at P5, P7, and P10. Bars plot mean + SE, number of cells in parentheses. P values by 2-way ANOVA adjusted for multiple comparisons. (C) Cumulative histogram of inter-event intervals for control and *Rorb* KO L4 barrel cortex at P5, P7, and P10.

### The putative RORβ target, Thsd7a, is required for adult TCA but not barrel wall organization

To begin exploring the relationship between disrupted gene expression in the *Rorb* KO and barrel organization, we examined known functions of genes differentially expressed at multiple developmental time points. Two candidates were identified with potential roles in cell migration and synaptogenesis. PlexinD1 (Plxnd1) is a cell signaling molecule known to play a role in pathfinding and synaptogenesis (Chauvet et al., 2007; Wang et al., 2015). Thrombospondin 7a (Thsd7a) regulates endothelial cell migration (Wang et al., 2010) but, it’s role in the brain is unknown. In controls, expression of both genes followed a similar developmental trajectory as RORβ, peaking around P7 (Figure 7A). In the *Rorb* KO, *Plxnd1* was significantly lower at P2 and P7 while *Thsd7a* was significantly lower at all three time points. In addition, we identified several differential ATAC peaks near *Thsd7a* with significantly reduced accessibility (Figure 7B). This included a peak containing a strong RORβ motif just downstream of the transcription start site, suggesting *Thsd7a* might be a direct target of RORβ regulation.

**Figure 7.**
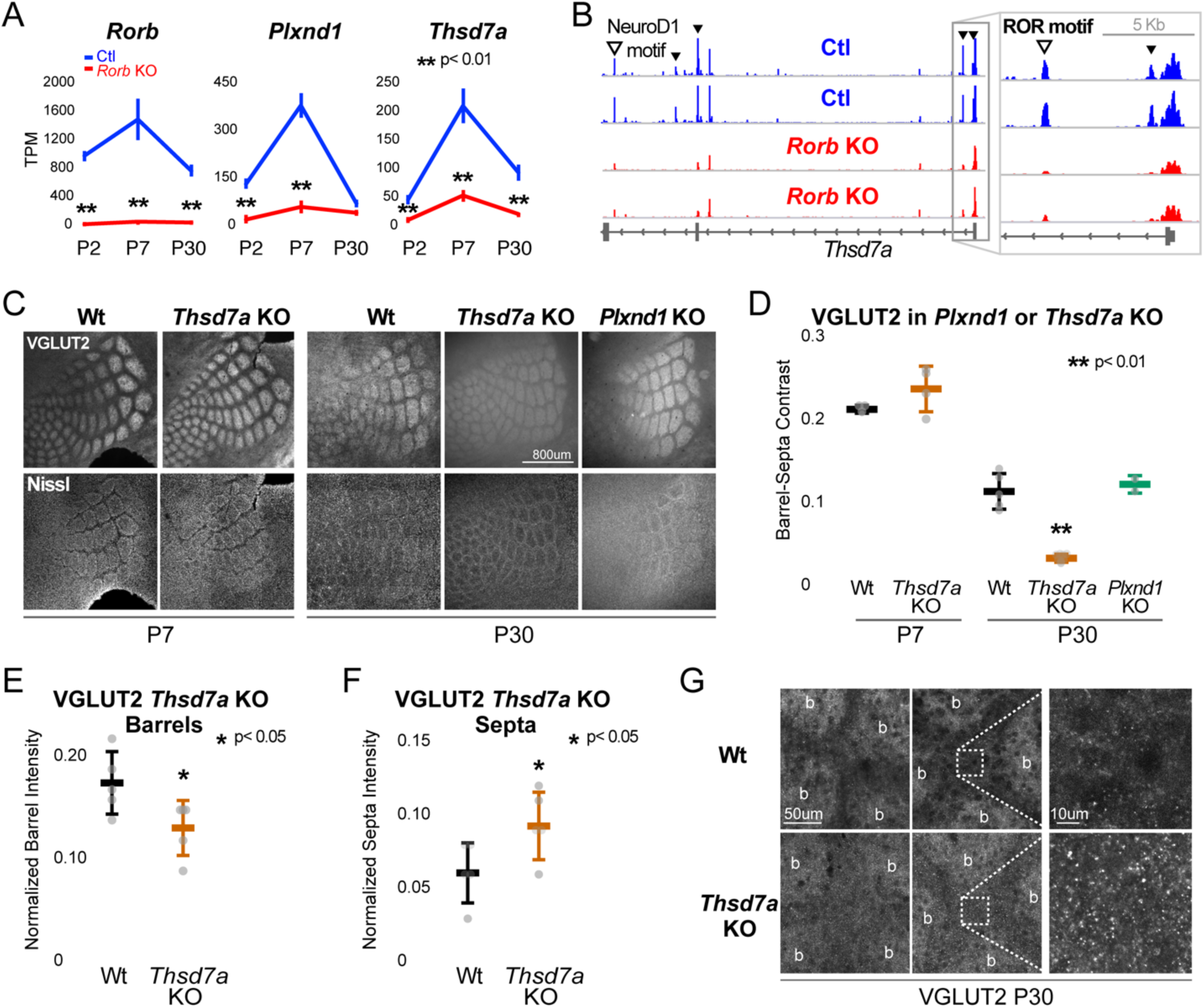
Thsd7a is required for TCA but not barrel wall organization. (A) Line plots of transcripts per million (TPM) measured by RNA-seq for three genes (*ROR*b, *Thsd7a*, and Plxnd1) from Ctl (blue) or *Rorb* KO (red) S1 layer IV barrel cortex. Lines plot the mean ± SE. (B) ATAC-seq around the *Thsd7a* gene (as in Figure 5A). (C) VGLUT2 and Nissl staining of whisker barrel cortex at P7 and P30 from wild-type (Wt), Plxnd1 KO, or Thsd7a KO. (D) Quantification of VGLUT2 Barrel-Septa Contrast from genetic lines in C. N=2-5 animals. Whisker plots as described for Figure 1B. (E) Background normalized quantification of VGLUT2 contrast in barrel hollows. Two tissue sections containing the largest portions of whisker barrel field were averaged per animal. N=5, P30 animals per genotype. Whisker plots as described for Figure 1B. (F) Background normalized quantification of VGLUT2 contrast in septa. Two tissue sections containing the largest portions of whisker barrel field were averaged per animal. N=5, P30 animals per genotype. Whisker plots as described for Figure 1B. (G) VGLUT2 staining imaged at high magnification (63X) in P30 Wt or *Thsd7a* KO whisker barrel cortex. Barrels are labeled “b”.

There was no detectable disruption to barrel organization in *Plxnd1* conditional KO mice (PlexinD1^flox^ crossed to Emx1-cre, Figure 7C-D). A *Thsd7a* constitutive KO also showed no disruption to barrel wall organization at P7 or P30. Interestingly, *Thsd7a* KO did show decreased VGLUT2 contrast between barrels and septa at P30 but not P7, suggesting Thsd7a is important for maintenance of TCA organization in adulthood (Figure 7C-D). The barrel phenotype of *Thsd7a* KO was qualitatively different from *Rorb* KO barrels. Specifically, the overall barrel pattern remained more intact in the *Thsd7a KO* despite the quantitative decrease in VGLUT2 contrast. *Thsd7a* KO may maintain sharper barrel borders than the *Rorb* KO due to intact barrel walls. Reduction in VGLUT2 contrast in the *Thsd7a* KO could be due to increased TCA localization in the septa and/or decreased TCA localization in the barrels. To distinguish these two possibilities, three regions of low VGLUT2 staining adjacent to the barrel field were quantified and used for within tissue slice normalization of barrel and septa intensities. *Thsd7a* KO resulted in a 24% decrease in barrel hollow VGLUT2 signal and a 56% increase in the septa (Figure 7E-F). High resolution imaging showed a clear increase in VGLUT2 puncta located in the septa (Figure 7G). Thus, loss of Thsd7a after *Rorb* KO likely contributes to the decrease in TCA segregation in adulthood.

## DISCUSSION

While somatotopic maps were one of the earliest and most obvious forms of cytoarchitecture, our understanding of the role neuronal identity plays in module formation is largely unknown. Studies have long approached the question of what drives cortical organization from the perspective of network activity and, in the case of barrel cortex from the perspective of key structures and pathways needed to relay sensory input. More recent studies characterizing transcription factors required in the cortex for barrel organization points to the importance of molecular mechanisms regulating transcriptional programs. However, many of these TFs are part of the pathways that carry sensory input or are fundamental regulators of broad developmental programs. It was unclear whether a TF such as RORβ, a highly restricted marker of L4 identity in the cortex, could influence macro-scale processes such as module formation. Indeed, we show that while RORβ is clearly regulating only a fraction of the phenotypic and transcriptional properties of L4 neurons, it is necessary for terminal specification of L4 identity and proper organization of L4 cytoarchitecture.

Specifically, RORβ is required in the cortex for barrel wall formation and full TCA segregation. This differs from earlier work focusing on the role of TCA patterning and activity as instructive for barrel wall formation. Instead, we find that loss of RORβ specifically in the cortex affects TCA segregation shortly after barrel walls should have formed, suggesting that bidirectional signaling between L4 neurons and TCAs is involved in establishing proper organization. Few other studies highlight this role of cortical influence on TCA organization (Iwasato et al., 2000).

While desegregation and loss of TCA patterning worsened with age, removing RORβ function after barrels form did not affect TCA segregation. From this we conclude the major contribution of RORβ occurs before and/or during barrel development. Once barrels have fully formed, RORβ activity is not required to maintain TCA segregation. We note, however, that loss of the putative RORβ gene target, Thsd7a, primarily affected TCA segregation in adults despite maximal expression at P7, which declines significantly by P30. One possibility involves Thsd7a functioning around the time of barrel formation to establish long lasting TCA structures that only manifest aberrant phenotypes later in life. Alternatively, the moderate expression level of Thsd7a at P30 may be sufficient for a role in adult maintenance. In either case, a role for Thsd7a in the nervous system has not been described previously. In endothelial cells, Thsd7a localizes to the membrane of the leading edge of migrating cells where it functions to slow or inhibit migration (Wang et al., 2010). Perhaps in somatosensory cortex it inhibits movement of nearby projections such as dendrites or axons allowing cortical neurons to “corral” TCAs in barrel hollows.

Our observation that barrel organization declined with age is very interesting and possibly the first description of this phenomenon in mice (Rice, 1985). It suggests continued plasticity or degradation of maintenance mechanisms over time. Few studies have examined plasticity within this structure in adulthood. This is in part because studies have shown a decline in the capacity to rewire sensory input to the cerebral cortex with age in certain systems. In the visual system, loss of sensory input has been shown to alter TCAs during a critical postnatal period (Antonini and Stryker, 1993; Erzurumlu and Gaspar, 2012). It is thought that once this critical period closes, TCA organization is fixed. Thus, developmental processes in the visual and somatosensory systems are assumed to stabilize TCAs and restrict learning and memory related changes to plasticity among cortical connections (Fox, 2002; Feldman and Brecht, 2005; De Paola et al., 2006; Karmarkar and Dan, 2006). However, there is some evidence to support a shift in this model of adult plasticity in both the visual and somatosensory cortex (Khibnik et al., 2010; Wimmer et al., 2010). In particular, Oberlaender et al. showed that a mild form of sensory deprivation induced by whisker trimming in 3-month old rats substantially altered TCAs in barrel cortex (Oberlaender et al., 2012). However, because adult TCA plasticity has garnered limited attention, we currently lack genetic studies examining the molecular mechanisms behind these processes. The natural decline in barrel organization and the mechanism of Thsd7a influence on TCA segregation merit further investigation as exciting new contexts to study both the functional roles of cortical organization and the impact of age.

Recent studies are revealing that neuronal identity in certain structures remains plastic during early postnatal periods. For example, mistargeted L4 neurons that migrate to layer 2/3 take on characteristics of their surroundings (Oishi et al., 2016b) and misexpression of some TF can alter the identity of postnatal neurons (Rouaux and Arlotta, 2010; 2013). We find that loss of RORβ shifts the transcriptional identity of L4 neurons to a more L5-like profile. This likely occurs through complex reorchestration of gene regulation. Upregulation of known L5 TFs such as Bcl11b/Citp and Etv1 at P2 may help drive an early diversion down an L5 trajectory. Regulatory signatures detected in adult neurons such as closure of binding sites for Zfp281 enriched near L4 genes and opening of Nfe2l1/3 motifs enriched near L5 genes may represent the tip of the developmental iceberg. In addition, our stringent motif analysis aimed to keep false positives low but may also miss relevant regulators with more minor roles. While we detect changes in binding capacity for many TFs, including RORβ, the complexity of dysregulation spread out across early postnatal development means there are certainly additional mechanisms driving this shift in cellular identity to be discovered. Here we combine the power of genetic knock-out strategies with multiple molecular profiling assays to interrogate the transcriptional network influenced by RORβ. We found RNA-seq paired with ATAC-seq provided a rich picture of the transcriptional changes occurring in *Rorb* KO neurons and insight into both developmental and adult functioning. Changes to the transcriptional network involved both differentially expressed TFs and TFs whose only perturbation was increased or decreased access to binding sites. Without these complementary perspectives, proteins such as Zfp281 and Nfe2l1/3 TFs might have been overlooked.

We identify several other TFs worthy of further investigation for their role in cortical development. Ascl1 and NeuroD1 are potent TFs that can induce transdifferentiation of mouse embryonic fibroblasts or microglia into neurons (Vierbuchen et al., 2010; Matsuda et al., 2019). NeuroD1 binds a different motif than NeuroD2, which is known to regulate barrel formation (Ince-Dunn et al., 2006), suggesting a distinct role. In addition, Trps1 was strongly upregulated by RORβ loss at P7 and P30, and was enriched in regions that opened. Its role in neurons is not clear, but it has been characterized as a transcriptional repressor that inhibits cell migration making it a tempting target to explore the lack of L4 neuron migration necessary to form barrel walls (Wang et al., 2018).

In addition to disrupted layer identity we also detect a significant disruption in the potential for *Rorb* KO cells to transcriptionally respond to activity connecting cellular identity, module formation and molecular responsiveness to input. In the adult *Rorb* KO, many activity-regulated TFs were upregulated, with the exception of Btbd3, and their DNA motifs showed increased accessibility. Around P7, when activity is critical for instructing cortical reorganization, we see reduced mEPSC frequency in L4 *Rorb* KO neurons, which is rectified by P10. Some of the transcriptional changes in the *Rorb* KO may be a form of compensation for the lack of input at P7. Failed upregulation of Htr1b and downregulation of S100a10/p11 may also be an attempt to increase activity in KO neurons. More is known about the role of Htr1b in TCAs where it is transiently expressed and, when stimulated, inhibits thalamic neuronal firing (Bennett-Clarke et al., 1993; Rhoades et al., 1994) and disrupts barrel formation (Young-Davies et al., 2000). TCA inhibition is thought to be the mechanism by which excess 5-HT disrupts barrels. While it is difficult to infer the role of Htr1b and p11 without characterizing cellular localization in S1 L4 neurons, downregulation of p11 resulting in less Htr1b localizing to the membrane coupled with reduced Htr1b expression could relieve inhibition in L4 *Rorb* KO neurons. Barrel formation and the ability to respond to activity inputs corresponds with increased RORβ expression and this increase is attenuated when TCA inputs are eliminated (Pouchelon et al., 2014). Together this suggests terminal differentiation and migration of neurons within L4 to form barrel walls are closely synchronized to excitatory input and require RORβ for proper establishment.

Although few other studies have examined the transcriptional targets and molecular mechanisms of TFs that regulate barrel formation, our study suggests RORβ is likely involved in the later stages of cellular specification and implicates several new TFs. RORβ also appears to function by distinct mechanisms from TFs previously characterized to regulate barrel formation. Loss of Bhlhe22 disrupts both barrel wall formation and TCA segregation but results in downregulation of Lmo4 (Joshi et al., 2008), unlike *Rorb* KO which increased Lmo4. Interestingly, Eomes is required for barrel wall organization but does not appear to affect TCA segregation (Elsen et al., 2013). Lhx2 and RORα are more broadly expressed than RORβ. *Lhx2* KO results in moderate down regulation of RORβ suggesting it is also likely upstream of RORβ in barrel development (Wang et al., 2017). Loss of Lhx2 greatly reduced TCA branching producing smaller barrels and barrel field. This phenotype is very similar to *Rora* KO barrels (Vitalis et al., 2017) suggesting RORα’s mechanism may be more similar to earlier developmental TFs than to RORβ. Disruption of barrelettes development in *Rora* KO thalamus is also consistent with a role in earlier stages of development. On the other hand, several TFs appear to be downstream of processes regulated by RORβ. For example, Btbd3 is important for dendritic orientation and is downregulated in the *Rorb* KO. It may be that dendritic orientation occurs after L4 cells have migrated to form barrel walls and provide an organized reference point for orientation. Thus, we have characterized in depth the molecular and transcriptional mechanism of RORβ as it orchestrates a critical juncture in barrel development where terminal differentiation and activity inputs are integrated to drive cellular organization in the cortex.

## Materials and Methods

### Animals

All animals were bred, housed, and cared for in Foster Biomedical Research Laboratory at Brandeis University (Waltham, MA, USA). Animals were provided with food and water *ad libitum* and kept on a 12hr:12hr light:dark cycle. Cages were enriched with huts, chew sticks, and tubes. All experiments were approved by the Institutional Animal Care and Use Committee of Brandeis University, Waltham, MA, USA.

*Rorb*^*GFP*^ (*Rorb*^*1g*^) and *Rorb*^*f/f*^ (*Rorb*^*flox/flox*^) mice were obtained from Dr. Douglas Forrest (Liu et al., 2013; Koch et al., 2017; Byun et al., 2019). *Rorb*^*GFP*^ mutation deletes the RORβ1 isoform, the predominant isoform in brain, and not the RORβ2 isoform (Liu et al., 2013). The *Rorb*^*f/f*^ allele deletes both isoforms. The following mice were obtained from Jackson Laboratories: TDTomato (stock 007909, RRID:IMSR_JAX:007909); plexinD1flox (stock 018319, RRID:IMSR_JAX:018319); Thrombospondin7a (Thsd7a) (stock 027218, RRID:MGI:6263683); EMX1-IRES-cre (Emx1-cre) (stock 005628, RRID:IMSR_JAX:005628); SertCre (Slc6a4) (stock 014554, RRID:IMSR_JAX:014554). CamK2a-cre (stock 005359, RRID:IMSR_JAX:005359).

### Perfusion

Animals were fatally anesthetized and transcardially perfused with 15mL 1x PBS (Fisher, SH3001304) then 15mL 4% PFA (Sigma Aldrich P6148-500G). Brains were fixed overnight in tangential orientation. After removing the whole brain from the skull, the cerebellum and olfactory bulbs were removed. The brain was split into two hemispheres along the longitudinal fissure and the midbrain was gently excised. The remaining cortex was placed in a shallow well made from a cryostat mold, filled with 4% PFA and a glass slide set on top for flattening. Brains were removed from PFA after 24-48 hours and stored in 30% sucrose/PBS solution at 4°C.

### Immunohistochemistry

50µm slices were made on a freezing Microtome (Leica SM 2010R). Controls and KOs were stained together in batches. Slices were permeabilized overnight at 4°C in 0.3% Triton-X100 (Sigma Aldrich, T8787) and 3% Bovine Serum Albumin (Sigma B4287-25G) in PBS. Slices were then incubated for 24 hours in primary antibody solution containing 0.3% Triton-X100 and 3% Bovine Serum Albumin in PBS at 4C. Primary antibody dilutions were as follows: Guinea pig anti-VGLUT2 (Millipore AB2251, RRID:AB_2665454) 1:500-1:1000, rabbit anti-VGLUT2 (Synaptic Systems 135 403, RRID:AB_887883) 1:250, chicken anti-GFP (Aves labs GFP-1020, RRID: AB_10000240) 1:500-1:1000. Slices were washed 3 times in PBS for 10 minutes each at room temp and then moved to secondary antibody solution containing 0.3% Triton-X100, 3% Bovine Serum Albumin, 10% normal goat serum. All secondaries were used at 1:500; Goat Anti-Rabbit Alexa Fluor 564 (Invitrogen A-11037, RRID:AB_2534095), Goat Anti-Chicken Alexa Fluor 488 (Invitrogen A-11039, RRID:AB_2534096), Goat Anti-Rabbit Alexa Fluor 633 (Invitrogen A-21070, RRID:AB_2535731), Goat Anti-Guinea Pig Alexa Fluor 647 (Invitrogen A-21450, RRID:AB_2735091). Slices were stained using Nissl (Invitrogen N21479) at 1:250 in PBS for 2 hours at room temperature, washed in PBS as before, and mounted in VECTASHIELD HardSet Mounting Medium (Vector Laboratories, H-1500, RRID:AB_2336787). Slides were stored at -20C and imaged within 1 week.

### Imaging and fluorescence quantification

Tissue was imaged on a Leica DMI 6000B Inverted Widefield Imaging Fluorescence Microscope or a Zeiss LSM 880 confocal microscope. All genotypes and age groups contained roughly even numbers of males and females. A minimum of two slices containing at least 5 intact barrels between rows B-D were quantified per animal. Experimenters were blinded to age and genotype during imaging and quantification. Regions of interest (ROIs) were drawn manually by a blinded researcher around 5-6 intact barrels from rows B, C, or D using Fiji (Schindelin et al., 2012). An ROI including the total space around selected barrels up to the edges of adjacent barrels was drawn to be used for calculating septa intensity (Figure 8). For *Thsd7a* KO and controls, three additional ROIs were drawn in the region adjacent to barrel cortex with low VGLUT2 signal to be used as background to normalize barrel and septa intensity. Custom MATLAB code was used to quantify the average fluorescence in ROIs. Septa intensity was calculated as septa total ROI intensity - sum(barrel ROIs). Contrast = (barrel - septa) / (barrel + septa). Contrast and normalized barrel and septa intensity were calculated and then averaged for 2 slices per animal. Two-way ANOVA was used to test for a significant effect of genotype and/or age as well as for an interaction between the two variables. Independent sample t-test was used to test for significant differences between genotypes at each age.

**Figure 8.**
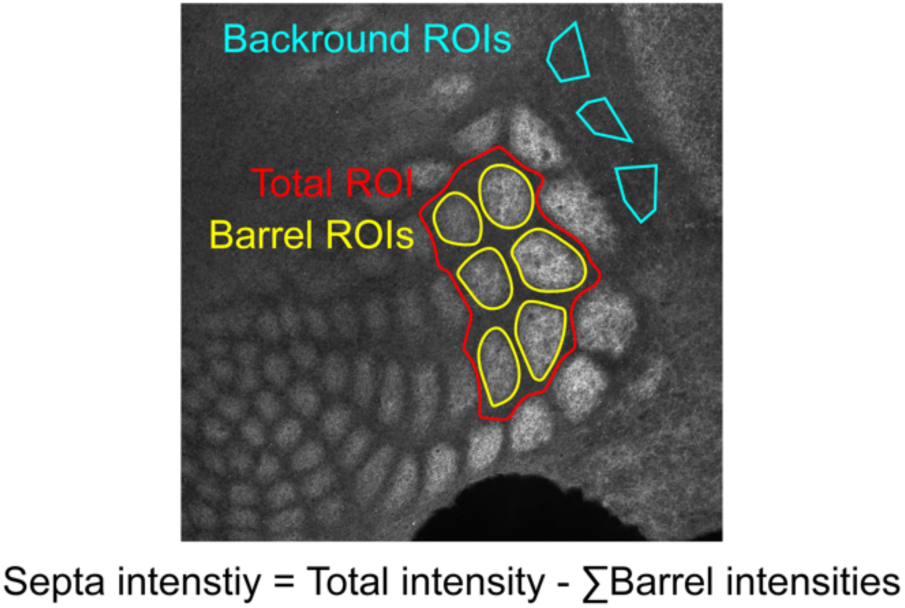
Example of quantification method. Regions of interest (ROIs) were drawn in Fiji by a researcher blinded to genotype and age.

### Electrophysiology

*Rorb*^*GFP/GFP*^ (KO) and *Rorb*^*GFP/+*^ (control; Ctl) mice were anesthetized with isoflurane and decapitated. Coronal slices (300µm) containing the primary somatosensory cortex were cut on a Leica (VT1000S) vibratome and incubated at room temperature in ACSF containing (mM) 126 NaCl, 25 NaHCO3, 2.5 KCl, 1.2 NaHPO4, 2 CaCl2, 1 MgCl2 and 32.6 dextrose adjusted to 326 mOsm, pH 7.4 and saturated with 95%/5% O2/CO2. Submerged, whole cell recordings were performed at 32 ± 1° on an upright microscope (Olympus BX50) equipped with epifluorescence. Pipettes with resistance 4-6 Mohm were filled with internal solution containing (mM) 100 K-gluconate, 20 KCl, 10 HEPES, 4 Mg-ATP, 0.3 Na-GTP, 10 Na-phosphocreatine and 0.2% biocytin adjusted to 300 mOsm, pH 7.35. For mIPSC recordings, the internal included 133 mM KCl and gluconate was omitted to bring E_Cl_ to 0mV. Recordings were made using an Axoclamp 700A amplifier, and were digitized at 10-20kHz and analyzed using custom software running under Igor 6.03 (Wavemetrics). Miniature synaptic events were recorded in voltage clamp at -70mV in the presence of PTX (mEPSCs) or DNQX+APV (mIPSCs) respectively.

Spiny stellate neurons were recognized based on their compact, GFP^+^ cell bodies within the GFP^+^ cell-dense layer 4. Input resistance was measured every 10-20 s with a small hyperpolarizing pulse and data were discarded if input or series resistance changed by >20%. P-values were calculated by 2-way ANOVA and adjusted for multiple comparisons by Tukey post hoc correction.

### RNA-seq

RNA-seq was performed as described previously (Sugino et al., 2019). Briefly, 1000-1500 GFP^+^ cells were isolated by FACS (BD FACSAria Flow Cytometer) from micro dissected L4 S1 live tissue (N=4 biological replicates per age and genotype). Figure 9 shows examples of the region micro dissected out to exclude L5. The four independent biological samples were collected from a pool generated by combining tissue from one male and one female mouse for a total of 8 animals used per time point. Cells were sorted directly into extration buffer and RNA stored at -80C for < three weeks. All libraries were prepared and sequenced in a single batch to prevent batch effects. Total RNA was purified (Arcturus PicoPure RNA Isolation kit, KIT0204) according to manufacturer’s specifications. Libraries were prepared using Ovation Trio RNA-Seq library preparation kit with mouse rRNA depletion (0507-32) according to manufacturer’s specifications and sequenced on a NextSeq Illumina platform (NextSeq 500/550 High Output (1 x 75 cycles)) obtaining 27 ± 2 million reads (mean ± SE). Reads were mapped by STAR with 90% ± 0.3% unique mapping (mean ± SE) and quantified with featureCounts (Liao et al., 2014). Differentially expressed genes were identified by Limma (Ritchie et al., 2015) using a fold change cutoff of 2 and padj<0.01 from a moderated t-test adjusted for multiple comparisons using FDR (Benjamini-Hochberg).

**Figure 9.**
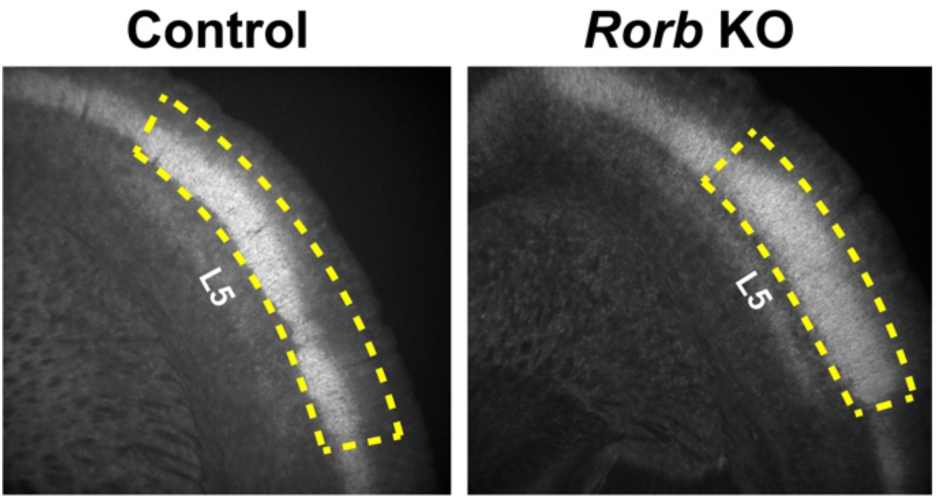
Example micro dissection of L4 S1 from coronal slices. Yellow dashed line indicates the outline of tissue retained for FACS.

### ATAC-seq

ATAC-seq was performed as described previously (Sugino et al., 2019). Briefly, 30,000 - 50,000 GFP^+^ cells were isolated by FACS from microdissected L4 live tissue (N=2 biological replicates per age and genotype). The two independent samples were collected from a pool generated by combining tissue from two male and two female mice for a total of 8 animals used. Nuclei were transposed for 30 minutes and libraries amplified according to published methods (Corces et al., 2017). Tagmented nuclei were stored at -20C for < two weeks. All ATAC libraries were purified, amplified, and sequenced as a single batch. Libraries were sequenced on a NextSeq Illumina platform (high output 300 cycles (2 x 150bp)) producing 105 ± 24 (mean ± SE) million reads per replicate. Reads were mapped using Bowtie2 and filtered producing 24 ± 2 (mean ± SE) million unique non-mitochondrial reads per replicate. TSS enrichment calculated per replicate according to the ENCODE quality metric (Corces et al., 2017) (https://github.com/ENCODE-DCC/atac-seq-pipeline) was 34 ± 3 (mean ± SE). Peaks were identified permissively using HOMER (-style dnase –fdr 0.5 -minDist 150 -tbp 0 -size 75 -regionRes 0.75 -region) (Heinz et al., 2010) and IDR (threshold=0.01, pooled_threshold=0.01) was used to identify reproducible peaks (Li et al., 2011). Differential ATAC peaks were identified using DiffBind with an FDR threshold=0.02 and log2 fold change in normalized read coverage threshold ≥1 (Ross-Innes et al., 2012).

RNA-seq and ATAC-seq datasets are available at GEO accession GSE138001.

### Motif Analysis

Motifs identified *de novo* from the sequences underlying ATAC peaks was carried out using MEME AME with shuffled input sequences as control and default settings (Fraction of maximum log-odds = 0.25, E-value threshold ≤ 10) (McLeay and Bailey, 2010), and HOMER findMotifsGenome.pl function masking repeats and -size given (Heinz et al., 2010). Scanning for specific motif matches in the sequences underlying ATAC peaks was carried out using MEME FIMO used the default threshold of p-value < 1e-4 (Grant et al., 2011) and HOMER findMotifsGenome.pl -find function. When possible 2-3 PWMs were obtained from Jaspar (Khan et al., 2018) and Cis-BP (Weirauch et al., 2014) prioritizing PWMs from direct data sources such as ChIP-seq. The R package GenomicRanges (Lawrence et al., 2013) was used to identify overlapping motifs between the two algorithms for cross validation. The overlap criteria allowed a 1 bp difference in the start or end position of the motif to accommodate ambiguity among motif models. Fisher Exact tests were calculated in R to test for enrichment of motifs in ATAC regions compared to control regions and to test for enrichment of genes with a nearby motif from a DEG group compared to a control group of genes. The set of control regions was generated by shuffling ATAC peaks throughout the genome excluding sequence gaps using BedTools (Quinlan and Hall, 2010) and the control group of genes were defined as expressed above 5 TPM but unchanged by age or *Rorb* KO.

## Acknowledgments

We thank Dr. Roland Schüle for agreeing to share the *Rorb*^*f/f*^ line, and Dr. Matthew Eaton for friendly bioinformatic advice. Supported in part by the intramural research program at NIDDK at the National Institutes of Health (DF).

## Supplemental Figures

**Figure 4 supplement.**
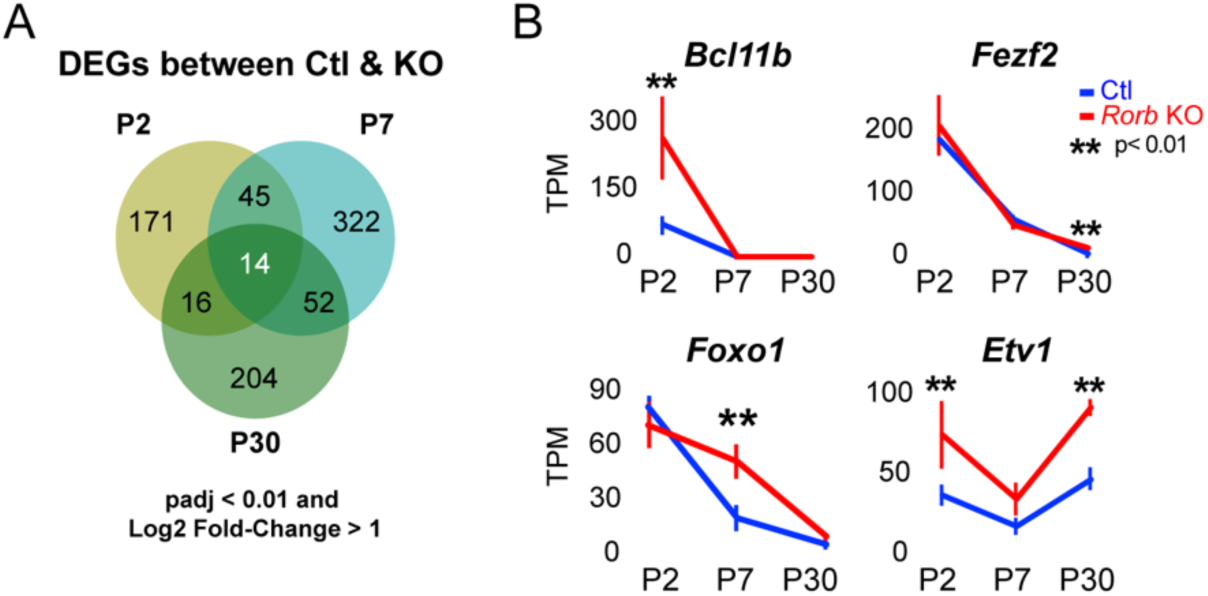
*Rorb* KO shifts the expression profile of neurons from a layer 4 to layer 5. (A) Differentially expressed genes (DEGs) at each age identified by RNA-seq. (B) RNA-seq expression of layer 5 TFs. Lines plot the mean ± SE. P by moderated t-test adjusted for multiple comparisons (Benjamini-Hochberg).

**Figure 5 supplement.**
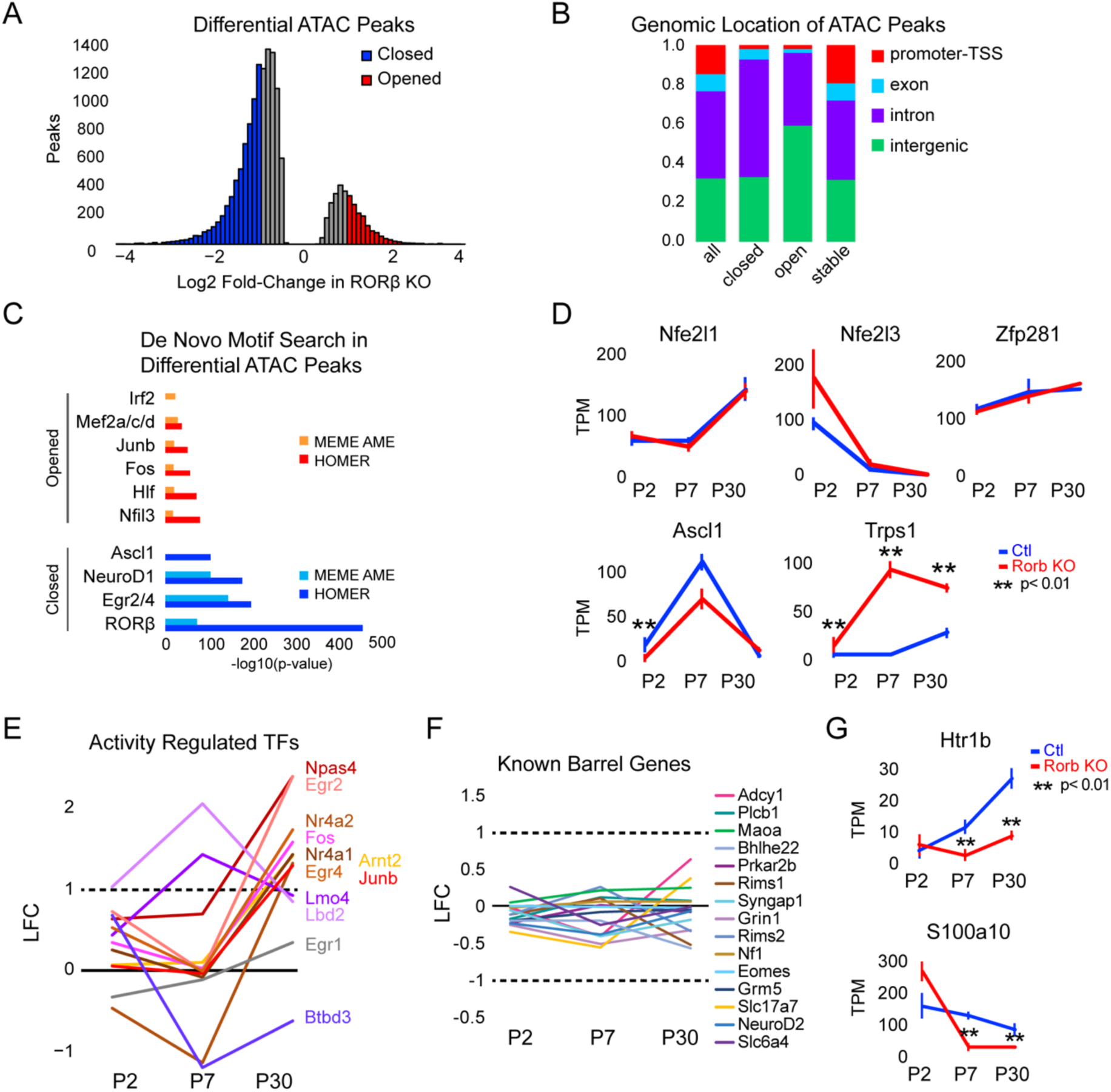
*Rorb* KO disrupts transcription factor binding sites near DEGs. (A) Differential ATAC peaks identified by DiffBind with Log2 Fold change (LFC) > 1 and FDR < 0.02. (B) Genomic distribution of ATAC-seq peaks identified in RORb control and *Rorb* KO. All peaks; all ATAC peaks in Ctl and KO, closed and opened peaks as defined in (B), stable; peaks with the lowest LFC between control and KO. Promoter defined as 2 Kb upstream of an annotated TSS. (C) De novo motif searching in differential ATAC peaks using two independent algorithms, MEME AME function and HOMER. Only motifs for TFs expressed in either sample are plotted. (D) RNA-seq expression of TFs with motifs in ATAC peaks. Lines plot the mean ± SE. P by moderated t-test adjusted for multiple comparisons (Benjamini-Hochberg). (E-F) RNA-seq log2 fold-c (LFC) for (E) activity-regulated transcription factors and (F) genes previously described to have a role in barrel organization. (G) RNA-seq of genes involved in serotonin signaling. Plots as in (D).

**Figure 6 supplement.**
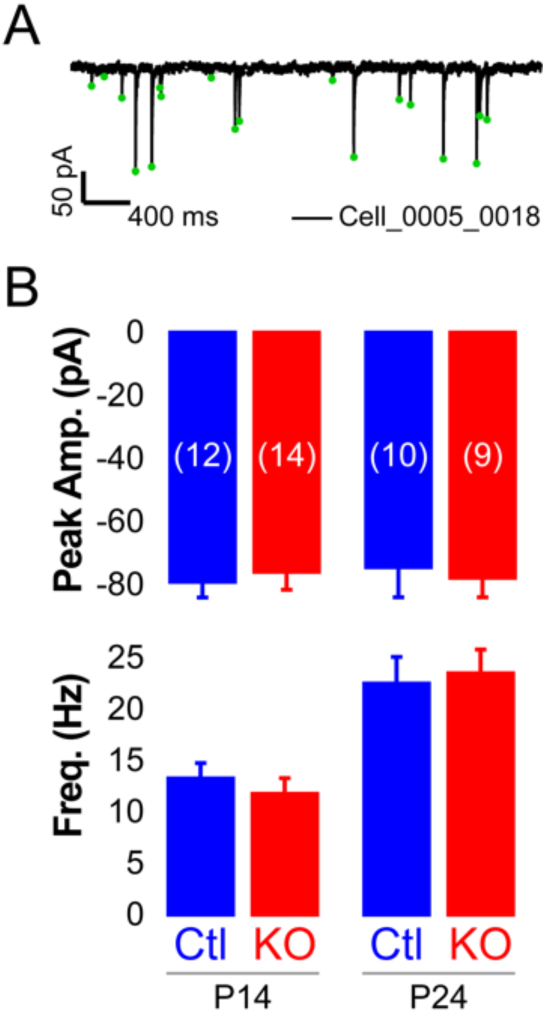
*Rorb* KO has minor effects on inhibitory input. (A) Example mini IPSC in one cell. (B) Mini IPSC amplitude (middle graph) and frequency (bottom graph) at P14 and P24. Bars plot mean + SE, N cells listed on upper graph.

